# Cell type matching across species using protein embeddings and transfer learning

**DOI:** 10.1101/2023.01.30.525343

**Authors:** Kirti Biharie, Lieke Michielsen, Marcel J.T. Reinders, Ahmed Mahfouz

## Abstract

**Motivation:** Knowing the relation between cell types is crucial for translating experimental results from mice to humans. Establishing cell type matches, however, is hindered by the biological differences between the species. A substantial amount of evolutionary information between genes that could be used to align the species is discarded by most of the current methods since they only use one-to-one orthologous genes. Some methods try to retain the information by explicitly including the relation between genes, however, not without caveats.

**Results:** In this work, we present a model to Transfer and Align Cell Types in Cross-Species analysis (TACTiCS). First, TACTiCS uses a natural language processing model to match genes using their protein sequences. Next, TACTiCS employs a neural network to classify cell types within a species. Afterwards, TACTiCS uses transfer learning to propagate cell type labels between species. We applied TACTiCS on scRNA-seq data of the primary motor cortex of human, mouse and marmoset. Our model can accurately match and align cell types on these datasets. Moreover, at a high resolution, our model outperforms the state-of-the-art method SAMap. Finally, we show that our gene matching method results in better matches than BLAST, both in our model and SAMap.

**Availability:** https://github.com/kbiharie/TACTiCS

## 1 Introduction

Model organisms, such as mouse and marmoset, are often used in brain research as a substitute for humans. However, because of differences between species, experiments performed on model organisms do not directly translate to humans. For example, widely-used antidepressants that target serotonin receptors are often tested on mice, while the expression pattern of serotonin receptors is highly divergent between human and mouse, likely leading to differences in cell function between species (Hodge *et al*., 2019). Consequently, to facilitate translational research, it is important to better characterize cell type matches between species. This facilitates studying how drugs then alter biological processes within specific cell types between these species.

Traditionally, cell types were characterized solely based on morphology, but using single-cell RNA sequencing (scRNA-seq), the expression pattern across thousands of genes can now be used to describe a cell type. This has resulted in the identification of an increasing number of cell types within specific brain regions (Tasic *et al*., 2018; Siletti *et al*., 2022). Although this improves our understanding of biological processes in the brain, when comparing species, it introduces the need for a method that can match these new cell types accurately between species.

Unfortunately, this is not a trivial task as genes are modified, duplicated and deleted throughout evolution, resulting in complicated many-to-many gene-gene relationships between species. These relationships become even more complicated when evolutionary distances increase.

Current methods that match cell types across species based on scRNA-seq data can be divided into two groups, mainly based on how they solve the gene-matching problem. The first group only uses the one-to-one orthologous genes, which are genes with exactly one match in the other species based on sequence similarity (e.g. using BLAST (Altschul *et al*., 1990)). Methods such as scANVI (Xu *et al*., 2021), MetaNeighbour (Crow *et al*., 2018), and LAMbDA (Johnson *et al*., 2019) belong to this group. While this is a straightforward approach, it ignores genes with a more complex evolutionary history which might have caused divergent functional specification of cell types between species. The second group of methods, including SAMap (Tarashansky *et al*., 2021), CAME (Liu *et al*., 2021), Kmermaid (Botvinnik *et al*., 2021), and C3 (Kabir *et al*., 2018), overcomes this limitation by considering many-to-many relationships between the genes based on sequence similarity. All these methods rely on the classical assumption that sequence similarity is a good measure of how genes functionally relate to each other. However, sequence similarity often considers one nucleotide/amino acid at a time, which to a large extent ignores sequence contexts important for functional characterization (e.g. secondary structures and sequence motifs). A growing body of evidence suggests that language models are a powerful approach to capture functional similarities between genes (Villegas-Morcillo *et al*., 2021; Rives *et al*., 2021; Heinzinger *et al*., 2019; Elnaggar *et al*., 2021). Similarly, we hypothesize that using language models to match genes between species can be beneficial for cell type matching.

Once we identified matching relationships between genes across species, the next step is to characterize cell type matches. We and others have posed cell type matching as a classification task where the agreement of predictions from two classifiers, trained on two labeled scRNA-seq datasets, is used to match cell types between the datasets (Michielsen *et al*., 2021; Johnson *et al*., 2019; Yuan *et al*., 2022). Biological differences between species, however, hinder applying such a method directly. A solution could be to learn a common embedding space for the cells before training the classifiers.

Here we introduce a method to Transfer and Align Cell Types in Cross-Species analysis (TACTiCS) that incorporates the two claims that we make: 1) using language models to match genes functionally between species, and 2) training classifiers in a shared embedding space to transfer cell types from one species to the other. We show that TACTiCS correctly matches human, mouse and marmoset brain cell populations from the primary motor (M1) cortex at a detailed cell type level, and does so better than SAMap, the current state-of-the-art method.

## 2 Methods

TACTiCS takes as input two single-cell (sc) or single-nucleus (sn) RNA-seq datasets, with raw expression counts, from two species A and B. TACTiCS consists of four steps (Fig. 1): 1) matching genes based on the protein sequences, 2) creating a shared feature space by mapping expression values with the gene matches obtained in step 1, 3) training within-species cell type classifiers, and 4) matching cell types by swapping the classifiers.

**Fig-1:**
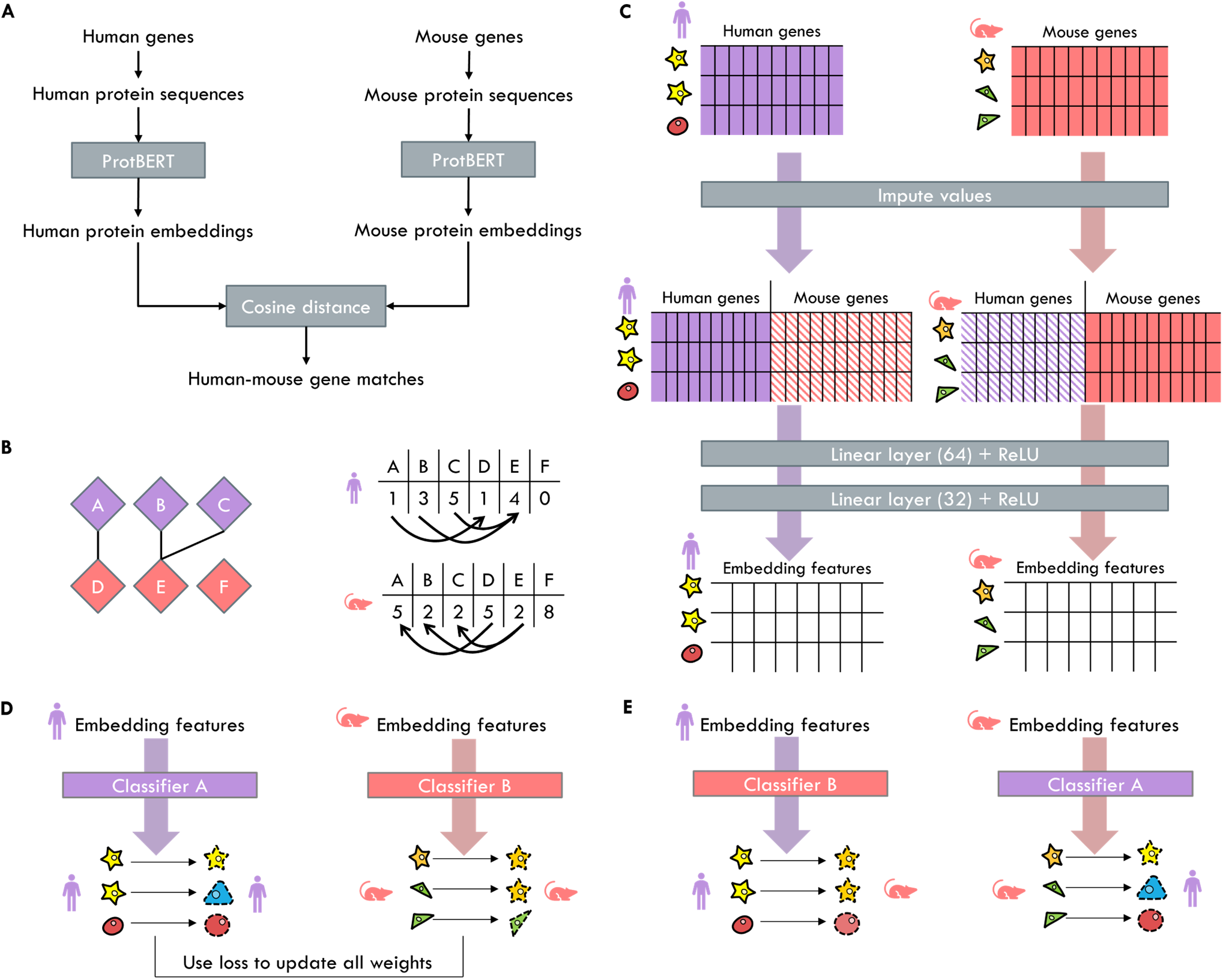
Schematic overview of TACTiCS. We use human and mouse as example, but cell types from any two species can be matched. (a) Matching genes on protein sequences using ProtBERT. (b) Bipartite graph of gene matches. Gene expression is imputed by taking the weighted average from connected genes in the bipartite graph. (c) Creating cell embeddings using linear layers on the shared feature space. The weights of the linear layers are shared. (d) Classifying within-species cells during training. The classifier consists of a linear layer outputting the cell type probabilities followed by a softmax. (e) Classifying cross-species cells using transfer learning. The predictions are used to match cell types.

### 2.1 Matching genes

First, we created an embedding for every gene using ProtBERT, a transformer-based language model (Elnaggar *et al*., 2021). The protein sequences were retrieved from UniProt (The UniProt Consortium *et al*., 2023). For human and mouse, we selected only the Swiss-prot sequences, but for marmoset we selected all protein sequences. We input the protein sequences to ProtBERT to create an embedding for each protein *h^ProtBERT^* (Fig. 1a). ProtBERT generates a 1024-dimensional embedding for every amino acid in the protein sequence. To allow TACTiCS to work with variable-length proteins, we followed common practice (Heinzinger *et al*., 2019) and took the mean embedding over all positions to represent the whole protein sequence (as well as the corresponding gene). Protein sequences longer than 2500 amino-acids (<2% of all sequences) were truncated to the first 2500 to fit into the memory of the GPU.

Next, for every pair of genes from species A and species B, we calculated the cosine distance between the ProtBERT embeddings. The initial set of gene matches were pairs with a cosine distance ≤ 0.05. To ensure that a gene is not connected to too many genes, we kept only the five closest genes, that met the distance threshold, for every gene.

Finally, we filtered the informative gene matches. Hereto, we calculated the top 2000 highly variable genes per species using Scanpy `highly_variable_genes`, and kept only those gene matches where at least one of the two genes is within the set of highly variable genes in their respective species (Wolf *et al*., 2018). From these matches, we construct two sets of genes *G_A_* and *G_B_*, corresponding to species A and B respectively, consisting of genes with a match in the other species.

To obtain sequence similarity-based gene matches, we used BLAST instead of ProtBERT to match the genes. We elected matches with an E-value <1e-6 as the initial set of matches. The bitscore of each match is used as the distance between the genes. Since BLAST scores are not symmetrical, one gene match is assigned a separate E-value and bitscore for each direction. If only one direction meets the E-value threshold, we use the corresponding bitscore as the gene distance. If both directions meet the threshold, we use the average of the two bitscores. The list of matches is then filtered similarly as before on that at least on of the genes in a pair is highly varying.

### 2.2 Creating a shared feature space by mapping expression values with the gene matches

We normalized the expression levels of genes as follows: 1) the raw expression counts of each dataset are normalized by the number of reads per cell such that the total number of counts in every cell is 10,000, and 2) the natural logarithm of the normalized counts are taken:

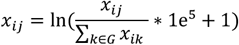

where *x_ij_* is the expression of gene *j* in cell *i*. Finally, a Z-score per gene is calculated to form the normalized expression matrices *X^A^* and *X^B^* for genes *G_A_* and *G_B_*, respectively. We created a shared feature space for the two datasets spanning *G_A_* ∪ *G_B_* (Fig. 1b). The shared feature space is partly equal to the expression matrices *X^A^* and *X^B^* and partly imputed:

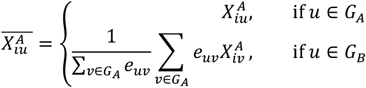

where 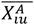 is the normalized expression of cell *i* from species A for gene *u* in the shared feature space. The expression of within species genes does not change. For a cross-species gene, we imputed the expression by taking the weighted average of the expression of the within-species genes it is matched to. The weight between gene *u* and gene *v* is calculated as:

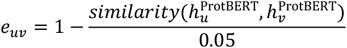

where *similarity* calculates the cosine distance between the ProtBERT embeddings. The weights are scaled to the interval [0, 1] by dividing with the distance threshold. When BLAST is used instead, we used the (average) bitscore between the two genes directly, since the bitscore does not have to be inversed. The edge weight is set to 0 for gene pairs that do not match according to the threshold and filtering criteria. The resulting matrices 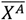 and 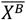 both span the same set of genes, and can thus be compared directly.

### 2.3 Cell embeddings

The shared feature space is put through two linear layers to create the cell embeddings (Fig. 1c). Each linear layer is followed by a Rectified Linear Unit (ReLU) activation function. The first layer creates embeddings of length 64. The second layer creates embeddings of length 32. These embeddings are used to visualize the embedding space with a UMAP. The weights to embed the cells are shared across the species.

### 2.4 Training species-specific cell type classifier

We used these embeddings to train a separate classifier per species. We used a neural network consisting of one linear layer followed by a softmax activation function (Fig. 1d). Both classifiers take the cell embedding as input and output cell type probabilities, *h^A,out^* or *h^B,out^*, only for cell types belonging to its respective species. During training, cells are input only to the classifier of its corresponding species.

The loss to update the embedding and classification weights consists of two parts: 1) the classification loss, and 2) the alignment loss. Both losses are calculated separately per species. For the classification loss, we used the weighted cross-entropy loss between the predictions and targets:

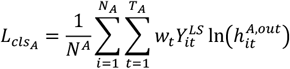

where *L_clSA_* is the classification loss for species A. *N_A_* and *T_A_* are the number of cells and cell types in species A respectively. *W_t_* is the weight for cell type *t*, explained further below. 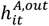 is the output of classifier A, specifically the probability that cell *i* belongs to cell type *t*. The one-hot encoded targets *Y* are modified with label smoothing to prevent overfitting and improve stability:

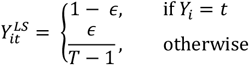

where *∈* (=0.1) controls the smoothness. The weight of each cell type is updated every epoch based on the accuracy of that cell type:

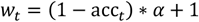

where *acc_t_* is the accuracy of class *t* in the current epoch. *α* is a hyperparameter that controls the influence of the accuracy on the weight. We use *α* = 9 such that the weights are in the interval [1,10] which restricts the relative difference in weight between cell types. By updating the weights, a cell type with a lower accuracy in the current epoch will have a higher weight in the next epoch and thus the predictions will shift to that cell type.

The alignment loss aims to integrate the embedding space across the species, such that cross-species cells with a similar gene expression are close in the embedding space:

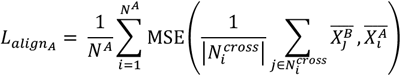

where *N^A^* is the number of cells of species A and 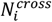 are the 20 nearest cross-species neighbours for cell *i*. MSE calculates the mean squared error between the prediction of the shared features of neighbours *j* and the actual shared features for cell *i*. If the alignment loss is minimized, neighbours in the embedding space can be used to predict the gene expression. The final loss is a combination of the classifier loss, the alignment loss and a regularization loss:

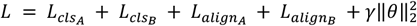

where *θ* consists of all parameters in the model, and is used for the L2 regularization to prevent overfitting. *γ* is the weight of the L2 norm, which is set to 0.01. The model is trained for 200 epochs. We used the Adam optimizer with a learning rate of 0.001. The full training process takes around 30 minutes.

To efficiently use large scRNA-seq datasets, the neural network is trained in batches. A batch size of 5000 cells per species is used to speed up the training while still having enough cells per cell type. Instead of sequentially iterating over the dataset, each batch is randomly sampled from the full dataset, while accounting for the size of each cell type. More specifically, every cell is assigned a probability 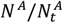 or 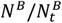, where *N^A^* is the total number of cells of species A and 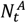 is the number of cells of species A belonging to cell type *t*. These probabilities are then used to sample a batch of cells per species with a similar number of cells for each cell type.

### 2.5 Transferring cell type predictions across species

After the neural network is trained, the cell types are transferred by using the classifiers on the species they were not trained on (Fig. 1e). That is, we calculate *h^B,out^* for cells of species A, and *h^A,out^* for cells of species B. The transferred cell type for a single cell is the cell type with the highest probability. To aggregate the information of the single cells to the cell type, we calculate the fraction of cells that are predicted to match cell types across species, which forms a normalized confusion matrix for both transferring directions. We average the two matrices to create a combined matrix, where high values indicate reciprocal matches. The values in the combined matrix can be used to score a match.

### 2.6 Dataset

We evaluated TACTiCS on snRNA-seq data taken from the primary motor cortex of human, mouse and marmoset (Bakken *et al*., 2021). These datasets consist of 76k human cells, 159k mouse cells and 69k marmoset cells, respectively. The cell type distribution varies considerably across species. For instance, non-neuronal cells make up around a third of both mouse and marmoset cells, while only 5% of the human cells are nonneuronal. We use two resolutions of the cell labels assigned by the original authors: 1) a higher resolution, consisting of 45 cell types present in all species; and 2) a lower resolution, consisting of 20 human, 23 mouse and 22 marmoset subclass cell types. At the lower resolution not all cell types occur in all species. SMC is only present in mouse, while Meis2 and Peri are only present in mouse and marmoset. Species-specific cells are labeled with “NA” at the higher resolution.

### 2.7 Evaluation

The combined matrix cannot be evaluated using standard metrics for confusion matrices, such as precision or F1 score, since we cannot distinguish between false positives and false negatives. Instead, we focus on the matching scores from corresponding cell types in the combined matrix, which ideally should be 1. We define the Average Diagonal Score (ADS) as the average score of the diagonal entries, after excluding species-specific cell types. A high ADS indicates that many cell types are correctly and reciprocally matched. However, the ADS does not indicate how many cell types are correctly matched. To this end, we define the recall as the fraction of diagonal entries where the score is highest for both that row and column.

We also compared TACTiCS to SAMap (Tarashansky *et al*., 2021), a cell type matching method that iterates between two steps. The first step matches the genes, which is initially done with BLAST on the DNA or protein sequences. Instead of taking the top-1 match, SAMap uses the BLAST bitscore directly in their model which allows for many-to-many matches. The second step uses the gene matches to first impute genes across species and then embed the cells by concatenating the principal components of the original expression and imputed expression. Then, the correlation between genes in the embedding space is used to update the gene matches. The two steps are repeated until the process converges.

### 2.8 Implementation

TACTiCS is implemented in Python 3.9. Pytorch (Paszke *et al*., 2019) was used for the model architecture. The scRNA-seq data is stored as Anndata (Virshup *et al*., 2021) objects, containing both the gene expression and the cell type annotations. The implementation of TACTiCS is available at https://2ithub.com/kbiharie/TACTiCS.

As Tarashanky et al. have noted, the runtime of SAMap increases significantly for larger datasets, and we were unable to run SAMap for the full datasets (Tarashansky *et al*., 2021). Instead, we used SAMap on subsets of 50k cells per species. We subsampled the data to keep the cell type proportions similar while making sure that all cell types are included.

## 3 Results

### 3.1 Matching genes using sequence embeddings is comparable to sequence alignment with notable differences

First, we investigate how similar the gene matches returned by ProtBERT and BLAST are. We retrieved 14,875 human and 13,601 mouse protein sequences, discarding 5% of the human genes and 13% of the mouse genes for which we do not have the protein sequence. We used both ProtBERT and BLAST to generate gene matches.

For 14,437 human genes, we found a mouse match using BLAST (gene with smallest E-value < 1e-6). For these human genes, we defined the ProtBERT match as the mouse gene with the most similar ProtBERT embedding. For 12,866 out of 14,437 human genes (89%), the BLAST match is identical to the ProtBERT match. Thus, the top-1 match is identical for the vast majority of genes. We ranked the BLAST matches according to the ProtBERT embedding distance to all mouse genes (Fig. 2a). Most of the BLAST matches have a rank close to 1 and over 95% of the BLAST matches have a rank below 100. Additionally, 31% of the BLAST matches that differ from the ProtBERT match are in the top-5 and thus considered in the many-to-many matches. Thus, if the BLAST match is not considered to be the best match by ProtBERT, it is still relatively similar based on the embedding distance.

**Fig. 2:**
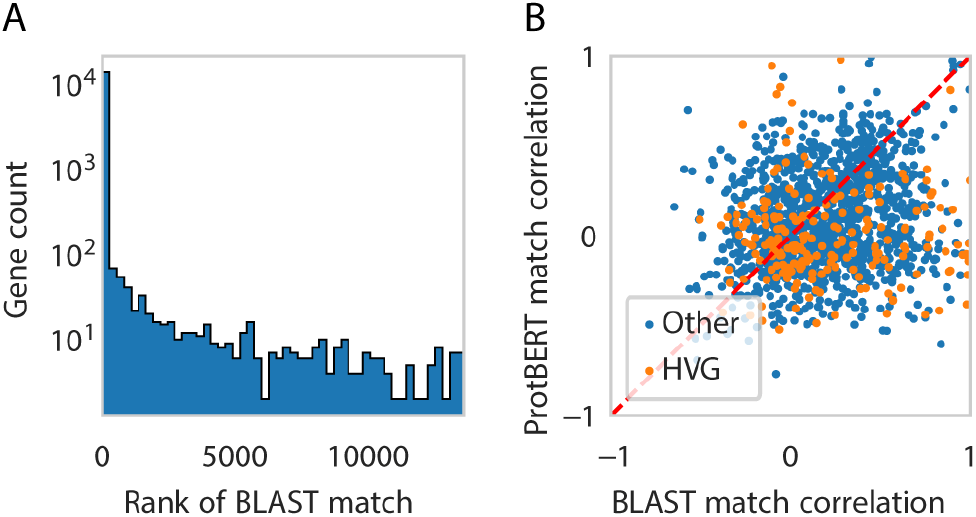
Comparison of ProtBERT and BLAST matches. (A) Rank of BLAST match according to ProtBERT embedding distances. Rank 1 indicates that the best ProtBERT match and the best BLAST match are the same. (B) Scatterplot of the correlation of the expression of human and mouse genes when considering the best BLAST match (x-axis) and the best ProtBERT match (y-axis). The expression correlation is calculated as the Pearson correlation across the average expression profiles of the Cross-species harmonized cell types. We omitted human genes where the BLAST match and ProtBERT match are the same. Gene matches where either the human gene, ProtBERT match or BLAST match is highly variable, are colored orange.

Next, we focus on the 1571 human genes for which the ProtBERT and BLAST match differ to investigate which method returns the most functionally similar match. We assess functional similarity here in terms of gene expression similarity across cell types. Therefore, we calculated the Pearson correlation coefficient across cell types in humans and mouse. We considered the harmonized cell types as defined in (Bakken *et al*., 2021) (Fig. 2b). For 1014 out of 1571 (65%) genes, the BLAST match has a higher gene correlation than the ProtBERT match. This is to be expected since the harmonized cell types were defined using the BLAST matches. However, for some genes, the ProtBERT match has a higher correlation than the BLAST match. For example, human *IL18R1* is matched to mouse *Il1r1* according to ProtBERT with a correlation coefficient of 0.945, while BLAST matches the gene to mouse *I118r1* with a correlation coefficient of 0.103 (Fig. 3). Human *IL18R1* and mouse *IlIrI* both show an increased expression for the endothelial and VLMC cells, while mouse *Il18r1* does not show this pattern, and is lowly expressed in all cell types.

**Fig. 3:**
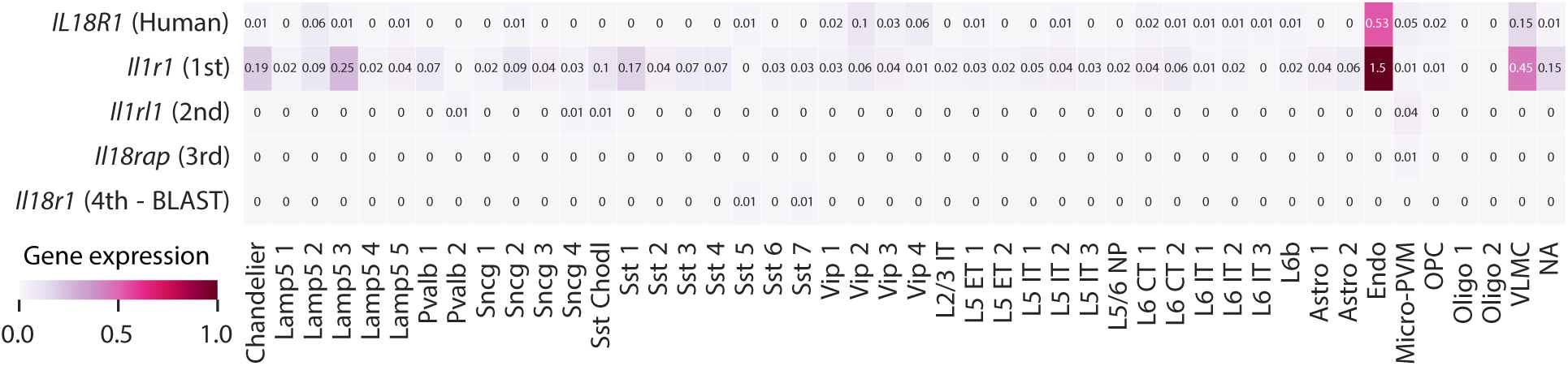
Average expression of human *IL18R1* and mouse matches across harmonized cell types. The mouse matches are ordered according to the ProtBERT embedding distances. BLAST matches human *IL18R1* to mouse *Il18r1*.

### 3.2 TACTiCS accurately matches cortical cell types across mouse and human

Now that we have seen that ProtBERT matches can be a powerful way to capture gene relationships, we use them in TACTiCS to match cell types in mouse and human cortex data. We use the Allen Brain Data, since the cell types have been carefully matched and harmonized by curators. We train TACTiCS for the human-mouse comparison for both the subclass and cross-species resolution. At the subclass resolution, TACTiCS returns the correct cell type for all 23 cell types that are present in both human and mouse (Fig. 4A). The species-specific cell types, mouse Meis2 and Peri, do not have a one-to-one match with a human cell type. For instance, mouse Meis2 matches human L6-IT and Vip with a low matching score in the combined matrix for both. Mouse Peri only matches human Sncg with a score of 0.5, but human Sncg matches mouse Sncg with a score of 0.8. Cell types present in both species have matching scores of ≥ 0.8 while wrong matches all have matching scores ≤ 0.6.

**Fig. 4:**
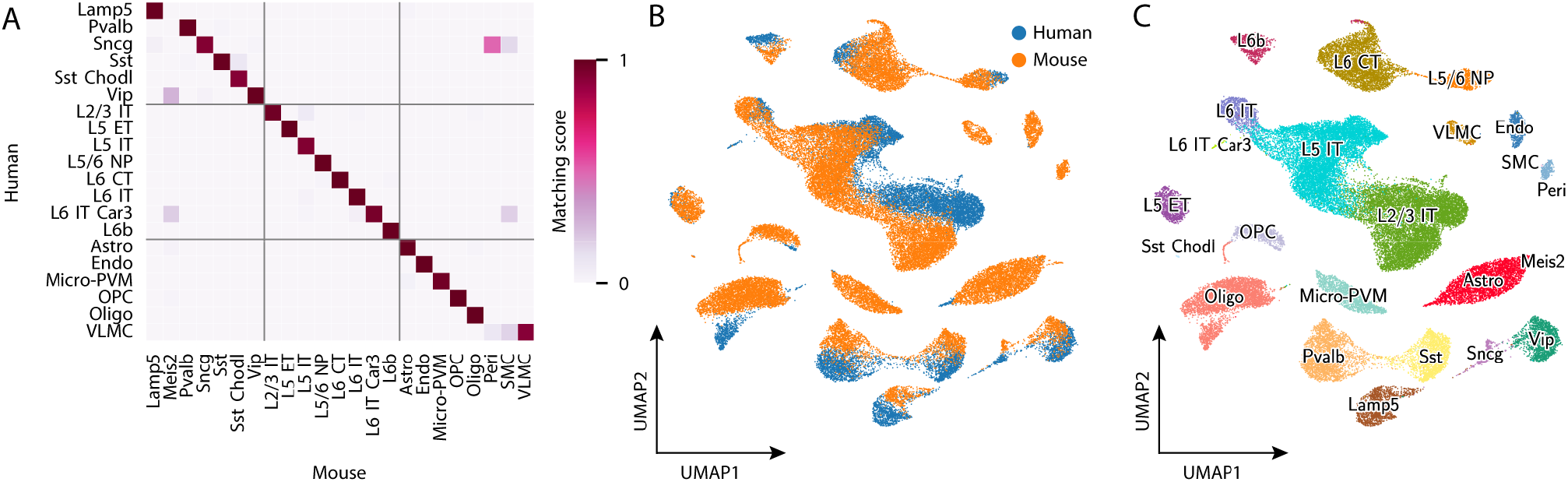
TACTiCS’ performance when matching human and mouse cell types at the subclass resolution. (A) Average confusion matrix of transferred cell types. (B) UMAP of cell embeddings, colored by species. (C) UMAP of cell embeddings, colored by cell type.

To get better insight into TACTiCS performance, we visualized the 32-dimensional cell embeddings using UMAP (Fig. 4b,c). Individual human and mouse cells do not mix well in the embedding space, but the UMAP does seem to align at the cell type level, i.e. corresponding cell types either overlap partially in the embedding space, or are relatively close. For example, Vip cells form a large cluster with partly human and mouse cells separated, and cells of mixed origin in the middle. The Sst cells also form a larger cluster, but the separation between the human and mouse cells is more visible. The Oligodendrocytes form two separate clusters, but they are closer to each other than to other cell types. The cell type proportions do seem to have an effect on the alignment in the embedding space. Cell types with a similar number of cells in human and mouse, such as Vip (6% in human and 2% in mouse), are clustered more coherently. Cell types with a large difference of occurrence within human and mouse, such as Astro (1% in human and 11% in mouse), cluster into one larger cluster with a smaller distinct but connected cluster for the species with the fewer number of cells. The mouse-specific cell types Meis2, Peri, and SMC are (correctly) clustered separately from the human cells. Thus, the embedding space can align the cell types across the species, but not the individual cells. Note that this can be due to unresolved batch effects or actual biological differences between the two species.

At the cross-species resolution, TACTiCS returns correct matches for the majority of cell types, with a recall of 0.89 (Fig. 5a, S1). Some cell types are not properly matched, namely the L5-IT subtype, some Lamp5 subtypes, some Sncg subtypes, and the Vip subtype. Thus, TACTiCS becomes relatively less accurate when there are a lot of subtypes that are not very distinct. In most cases, when there is a mismatch, TACTiCS matches a cell type with a subtype that belongs to the same subclass. The human Sncg subtypes are an exception since some of its subtypes are matched to mouse L5 IT or Lamp5 subtypes.

**Fig. 5:**
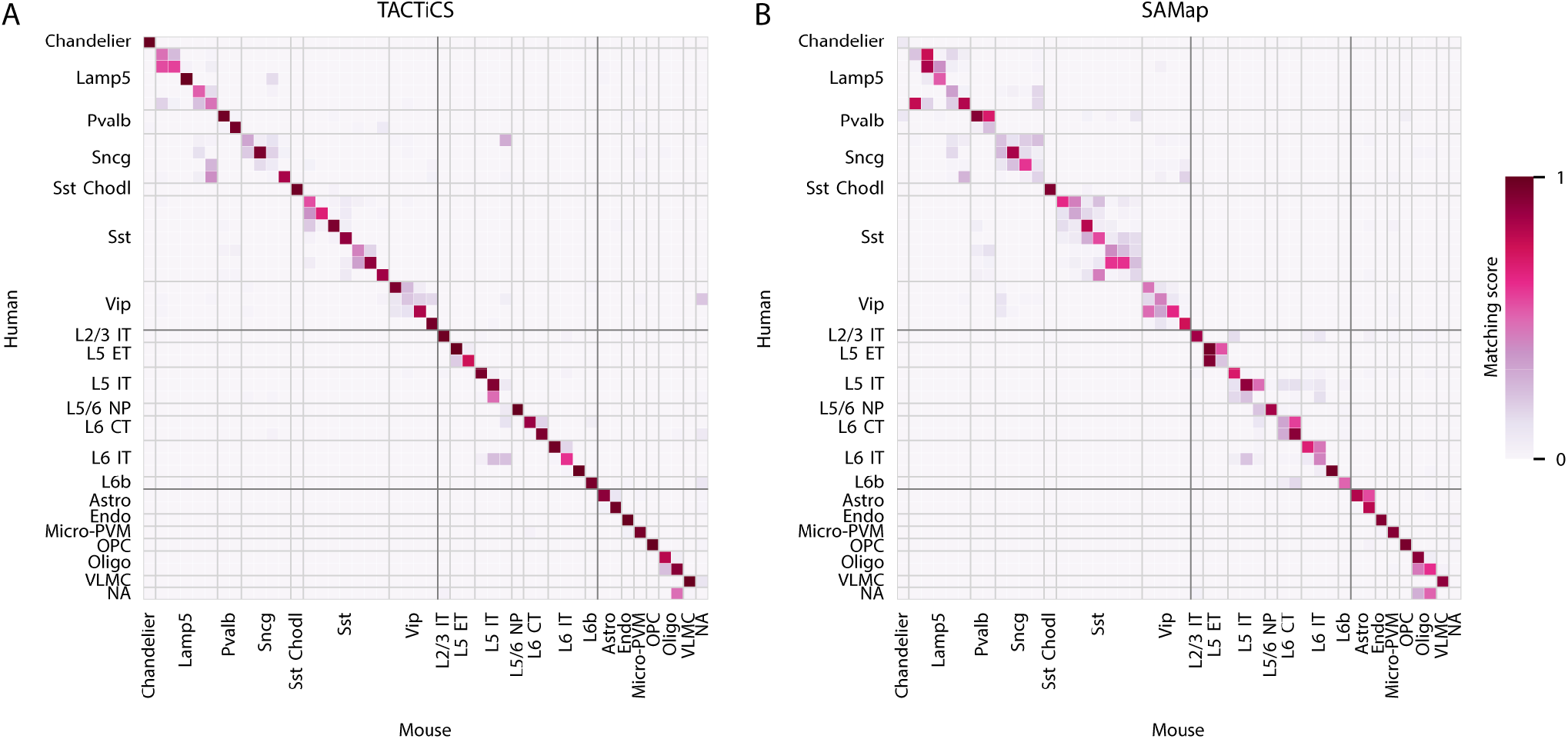
Performance of (A) TACTiCS and (B) SAMap on when matching human and mouse cell types at cross-species resolution. Cross-species cell types are grouped per subclass (indicated with the light-grey lines) and class (indicated with dark-grey lines).

To evaluate the performance of TACTiCS across species with variable evolutionary distance, we tested TACTiCS on cortical cell types between human-marmoset and mouse-marmoset (Table 1). At the subclass resolution, TACTiCS performs similar on all three comparisons with a recall of 1.0. At the cross-species resolution, TACTiCS performs best for the human-marmoset comparison and worst for the mouse-marmoset comparison. These results indicate that the performance of TACTiCS is dependent on the evolutionary distance between the species, since the evolutionary distance to the closest common ancestors from human and marmoset (~40mya) is a lot less than human and mouse (~70mya).

**Table 1:** Average Diagonal Score (ADS) and recall for TACTiCS and SAMap on human, mouse and marmoset. The gene-gene matching is either done using ProtBERT (P) or BLAST (B).

| Comp | Meth | Matching | Subclass |  | Cross-species |  |
| --- | --- | --- | --- | --- | --- | --- |
|  |  |  | ADS | Recall | ADS | Recall |
| Hu-mo | TACTiCS | P | 0.976 | 1.0 | 0.794 | 0.911 |
| Hu-mo | SAMap | B | 0.854 | 1.0 | 0.584 | 0.6 |
| Hu-ma | TACTiCS | P | 0.968 | 1.0 | 0.905 | 0.956 |
| Hu-ma | SAMap | B | 0.925 | 1.0 | 0.776 | 0.889 |
| Mo-ma | TACTiCS | P | 0.975 | 1.0 | 0.682 | 0.756 |
| Mo-ma | SAMap | B | 0.839 | 0.909 | 0.595 | 0.578 |
| Hu-mo | TACTiCS | B | 0.840 | 0.85 | 0.506 | 0.578 |
| Hu-mo | SAMap | P | 0.855 | 1.0 | 0.597 | 0.644 |

### 3.3 TACTiCS outperforms SAMap in matching cortical cell types across mouse, human, and marmoset

To benchmark TACTiCS, we compare its performance to SAMap using three pair-wise comparisons (human-mouse, human-marmoset, and mouse-marmoset). Across all comparisons, TACTiCS has a higher ADS and recall than SAMap (Table 1). Both methods perform well at the sub-class resolution for all comparisons with a recall of 1.0. However, TACTiCS assigns higher scores to correct matches than SAMap. For instance, SAMap correctly matches human L6b to mouse L6b, but with a very low matching score equal to 0.35, while TACTiCS matches the same cell types with a matching score of 0.99. Interestingly, for the species-specific cell types, TACTiCS suggests matches that have a low score (0.18-0.68), allowing to detect the species-specific cell types. The performance of SAMap for the species-specific cell types is not consistent across all cell types and comparisons. For example, SAMap correctly assigns zero scores to mouse Peri and SMC in the human-mouse comparison, but incorrectly matches mouse SMC to marmoset Peri with a high matching score.

At the cross-species resolution the performance of both methods drops compared to the subclass level as expected, but the difference between the methods becomes more apparent (Fig. 5, S1). For mismatches between subtypes, TACTiCS usually matches to subtypes within the same subclass, while SAMap regularly maps to cell types from another subclass. While both TACTiCS and SAMap partly match human Sncg to mouse Lamp5, SAMap additionally shows similarity between human Sncg and mouse Vip. SAMap also mismatches mouse Chandelier with human Pvalb, while TACTiCS matches human and mouse Chandelier with high confidence.

### 3.4 Using ProtBERT matches improves the cell type matching for both TACTiCS and SAMap

Finally, we assessed the importance of using the ProtBERT embeddings to match genes compared to using BLAST on the final cell type matches. To this end, we trained TACTiCS based on the BLAST matches and SAMap using the ProtBERT matches on the human-mouse data. For a fair comparison of ProtBERT to BLAST in SAMap, we only apply the embedding distance threshold to the ProtBERT matches, rather than filtering the gene matches thoroughly. Training TACTiCS at the cross-species resolution using the BLAST matches results in an ADS of 0.54 and a recall of 0.56, which is a lot worse than using the ProtBERT matches (Table 1). For SAMap, the ADS increased from 0.58 to 0.60 and the recall increased from 0.60 to 0.64 (Table 1) when ProtBERT matches were used instead of the BLAST matches. Thus, for both TACTiCS and SAMap it is beneficial to use ProtBERT embeddings to match genes.

## 4 Discussion

Here, we present TACTiCS, a method to accurately match cell types from scRNA-seq data across species. We applied TACTiCS to match cell types across human, marmoset, and mouse motor cortex, species with different evolutionary distances to each other. Even though TACTiCS matches cell types from all three species with high confidence, we showed that human and marmoset cell types are considerably easier to match which correlates with their closer evolutionary distance. Furthermore, we showed that TACTiCS outperforms the state-of-the-art method SAMap on all comparisons with the biggest difference at a higher resolution in favor of TACTiCS. We should note that here we focus on comparisons across species with a relatively small evolutionary distance while SAMap was originally developed to match cell types across larger evolutionary distances (Tarashansky *et al*., 2021).

Even though TACTiCS outperforms SAMap on the (finer) cross-species resolution, its performance drops as well. We would like to note that the cell types at this resolution were established by Bakken et al. by integrating datasets from the different species and clustering them in an embedding space (Bakken *et al*., 2021). This resulted in ambiguous clusters which were resolved manually by the authors to determine which cell types would be in one cross-species group. Since these matches are not perfect, it makes sense that we cannot achieve a perfect performance either. Furthermore, the ground-truth matches used for evaluation are based on analyses performed using BLAST one-to-one matches, also causing unwanted differences when comparing results.

Gene matching is one of the main components of TACTiCS. We match genes based on the distance between their corresponding protein embeddings, which are generated using ProtBERT instead of the commonly used sequence similarity based on BLAST. Even though the top-1 matches of ProtBERT and BLAST are largely similar, we have shown that using Prot-BERT instead of BLAST distances improves the performance of both TACTiCS and SAMap. When aligning sequences using BLAST, every amino acid is considered to be equally important, while we speculate that ProtBERT focuses more on functional domains. A downside, however, of using ProtBERT distances is that the protein sequence is needed and as a consequence, we can only use coding genes. Using DNA sequence embedding models, e.g. DNABert (Ji *et al*., 2021), for non-coding genes, could in the future be used to overcome this limitation.

Some cell types, such as Meis2 and Peri in mice, are species-specific. A limitation of our current approach is that the classifiers we built in TACTiCS are missing a rejection option and therefore we cannot identify these species-specific cells automatically. Since TACTiCS usually assigns a low matching score to these cell types, this score could be used to filter species-specific cells. This difference in scores, however, is mainly apparent at the subclass resolution. At the cross-species resolution, it would be more difficult to determine a proper threshold without deteriorating the performance. In general, SAMap assigns lower scores to species-specific cell types, but also for SAMap the difference is smaller at the cross-species resolution.

When inspecting the cell embeddings in the low dimensional space, we notice that the cells from difference species are not well mixed. Matching cell types, however, are closest to each other and species-specific cell types are more separated from all other cells. There are many data integration methods developed for single-cell data, such as scVI and Seurat (Lopez *et al*., 2018; Hao *et al*., 2021), that would achieve a significantly better integration. Since data integration is not the main goal of TACTiCS, we did not add an explicit mixing component to the loss function. The current loss function enforces that neighboring cells from the other species can predict the other cell’s gene expression profile. This enforces cells of the same cell type to be the closest, but not to fully overlap. Adding a component to the loss that forces cells to be mixed (e.g. to have neighbors of both species) could greatly improve the integration. Alternatively, if good integration is a user’s desire, an option would be to replace the component of TACTiCS that generates the cell embeddings with another data integration method such as scVI. The flexible architecture of TACTiCS allows the individual components (gene matching, cell embedding, and cell classification) to be easily replaced, extended, or integrated with different methods.

With TACTiCS we showed that using protein embeddings to match genes is a viable alternative to BLAST when matching cell types based on their scRNA expression levels across species. TACTiCS can accurately match cell types at different resolutions for large datasets, outperforming SAMap. We envision that this fast and accurate cell type matching method, will make comparative analyses across species considerably easier, contributing to, e.g. to the study of cell type evolution or translational research.

