## Supplementary Figures for "Cell type matching across species using protein embeddings and transfer learning"

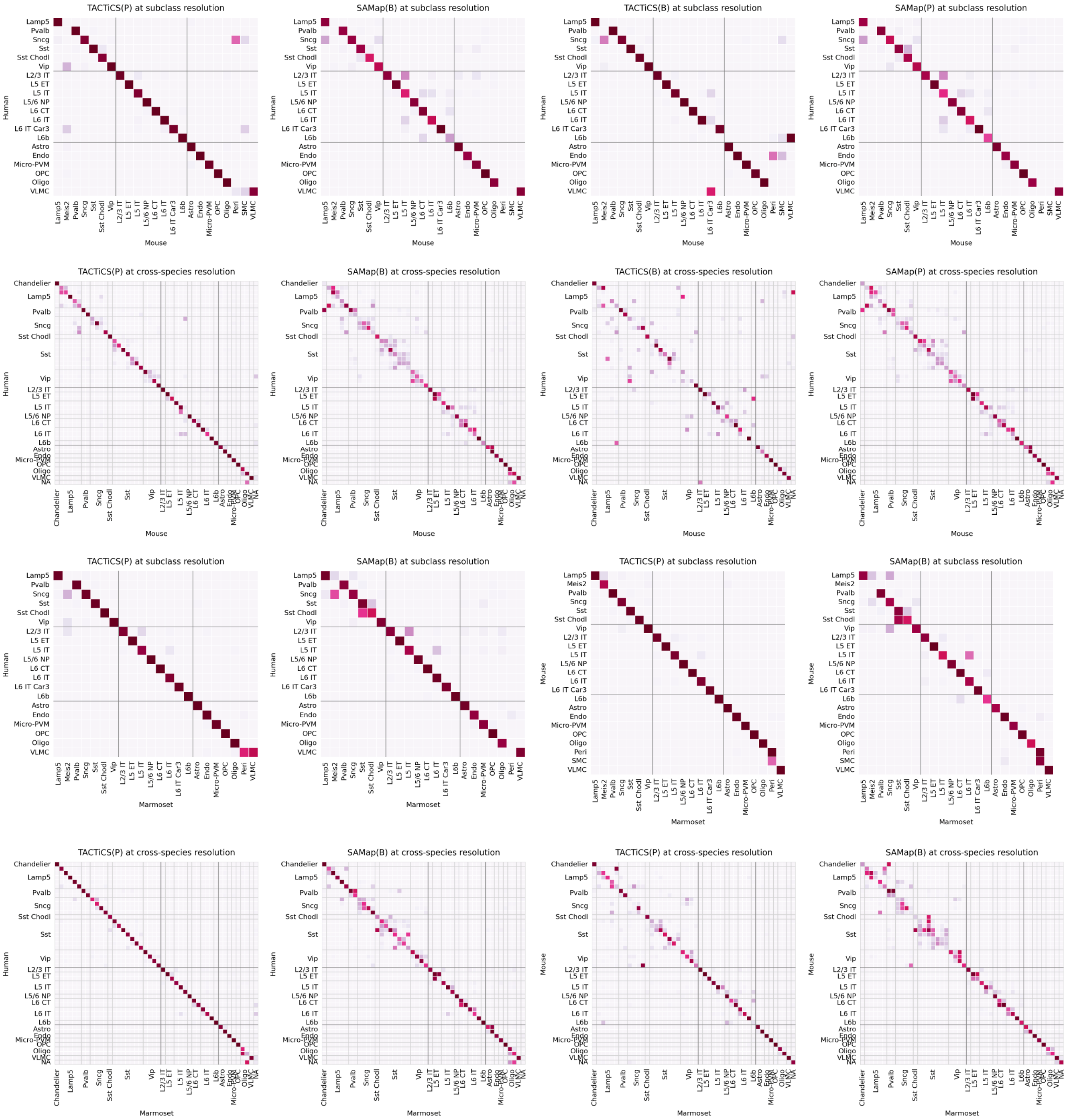


**Fig. S1:** Performance of TACTiCS and SAMap on three pairwise comparisons (human-mouse, human-marmoset, and mouse-marmoset) using ProtBERT (P) or BLAST (B) as distance metric on the subclass and cross-species resolution.
